# Motile and matrix-producing cells drive distinct modes of cell-scale motion in *Bacillus subtilis* colonies

**DOI:** 10.64898/2026.05.21.727008

**Authors:** Jason Samaroo, Abdulrahmen Almodaimegh, Joseph W. Larkin

## Abstract

The physics of cell motion and growth drives the expansion of groups of cells including bacterial colonies. Under different conditions, many species of bacteria grow with strikingly different dynamics, which can arise from the physics of constituent cells. For example, on low-percentage agar substrates, colonies often expand rapidly via motility as swarms, while on higher percentage substrates, biofilms expand slowly due to growth and pressure. However, in bacterial species *Bacillus subtilis*, both swarm and biofilm phenotypes contain differentiated cells in both the motile state and the matrix-producing state, provoking the question: how does phenotypic heterogeneity influence the cell-level motions that drive development? To answer this question, we used a bead-tracking assay in which we follow the motion of cell-sized fluorescent beads embedded in growing *B. subtilis* colonies. By being pushed due to their interactions with cells, beads passively report on local cell motion. We found that, in rapidly expanding swarms, bead motion was bimodal: some beads moved rapidly and diffusively, while others appeared to be slow or stationary. However, after following the seemingly trapped beads over many hours, we found that the beads in fact moved slowly and ballistically. In biofilms grown on higher percentage agar, we found that some beads were trapped, moving sub-diffusively over the course of biofilm growth, while others moved ballistically. We hypothesized that slow, ballistic bead motion in both swarms and biofilms was driven by groups of radially expanding matrix-producing cells. To test this hypothesis, we performed bead tracking experiments in colonies formed by regulatory mutant strains that were locked into motility or matrix production. In swarms of motile-only cells, we observed fast, diffusive bead motion with no population of slow, ballistic beads. In matrix-only biofilms, we observed that ballistic motion was heavily favored compared to wild type. These results support the hypothesis that expanding clusters of matrix cells drive slow, ballistic motion at the cellular scale during colony growth. Our results demonstrate that heterogeneous cell phenotypes contribute to heterogeneous local physics in bacterial colonies, influencing the distributions of cellular phenotypes.

## Introduction

Bacterial colonies expand through forces arising from cellular growth and physical phenomena such as cellular motility and extracellular matrix (ECM) production. Under different environmental conditions, colonies of genetically identical bacteria have been observed to expand with dramatically different phenotypes. For example, many species expand rapidly as a swarm of motile cells on low-percentage agar substrates and as extracellular matrix (ECM)-rich biofilms on high-percentage agar substrates^1–7^. These two modes of expansion have often been modeled with distinct physical mechanisms at the colony level. Swarms are generally treated as spreading fluids and biofilms are often modeled as growing viscoelastic solids^8–14^. These models can account for multiple aspects of colony growth such as surface morphology and the temporal dynamics of expansion.^10–12,14–17^ At the cellular level, swarms are assumed to be populated by flagellated motile cells and biofilms are assumed to be populated by matrix-producing cells, assumptions generally supported by gene expression data.^18^ However, cellular phenotypes in both kinds of colonies are often mixed. Biofilms contain cells expressing flagellar genes and swarms contain cells that synthesize matrix.^3,19^ These observations suggest that the internal physics of swarms and biofilms may be heterogeneous, however no study has examined how the internal, cell-level physics of swarms and biofilms depend on the cellular phenotypic composition of the colonies.

The question of how cell phenotypes behave within colonies is especially relevant in *Bacillus subtilis*, which is a model species for both swarms and biofilms^6,20^. Motility and matrix production genes are regulated in such a way that motile and matrix cell types are mutually exclusive^21,22^. In biofilms grown on 1.5% agar with MSgg medium, both matrix cells and motile cells are present, with matrix cells dominating the expanding exterior.^3,6,23^ Swarms grown on 0.5% agar also contain cells in both the motile and matrix states^24^. It is not known how the motile and matrix cell phenotypes themselves contribute to the internal physics of expanding colonies. Answering this question has been hindered by the inability to control cell phenotypes and measure motion over the large temporal and spatial scales of growth within a colony.

Here we measure cell-level motion generated during growth of *Bacillus subtilis* colonies using a bead assay in which we track the trajectories of 1 µm fluorescent beads embedded in colonies. We image swarms grown on 0.5% agar and biofilms grown on 1.5% agar. Both colony types contain cells expressing both flagellar genes and matrix production genes. We find that in swarms beads exhibit a dichotomy in motion. Some beads move rapidly with Brownian-like motion, while others move orders of magnitude more slowly with directed, ballistic motion. In 1.5% agar biofilms, beads exhibit a broad range of motions from sub-diffusive to ballistic. To isolate the effect of motile cells and matrix-producing cells on the internal dynamics of expansion, we perform experiments with regulatory mutants that are locked into motile phenotypes and matrix-producing phenotypes. We find that motile cells are associated with a distribution of fast and slow motions resulting in diffusive and ballistic motion occurring locally. Matrix producing cells were associated with ballistic motion within colonies. Our results demonstrate that heterogeneity at the cellular level creates heterogeneous dynamics within growing colonies that regulate internal cellular motion and phenotypic distributions.

## Results

### *B. subtilis* colonies show distinct colony expansion dynamics and phenotypic composition during growth in different environments

*B. subtilis* colony expansion and cellular phenotypic composition change with environmental conditions, for example across substrates of different agar percentage. To examine the effect of substrate stiffness on *B. subtilis* colony expansion and cellular phenotypic composition, we grew colonies of the minimally domesticated strain NCIB3610 (3610 hereafter) on MSgg (Minimal Salts glycerol glutamate, see Materials and Methods) substrates with 1.5% and 0.5% agar. As reported in previous works, 3610 grew as a wrinkled biofilm on 1.5% agar ^25^ (Fig. 1A) and as a motile swarm on 0.5% agar^26^ (Fig. 1B). Swarms expanded radially much faster than biofilms (Fig. 1C,D) as quantified in the radius vs time graph of Fig. 1E. Biofilms are thought to be characterized by extracellular matrix producing cells, while swarms are thought to be dominated by hyperflagellated motile cells.^27^ Consequently, the expansion of biofilms has been modeled as dominated by the physics of matrix production, while the expansion of swarms has been modeled as dominated by swimming cells.^12^ In *B. subtilis* cells, matrix production and motility are governed by a regulatory network that causes the two phenotypes to be mutually exclusive.^28,29^ Moreover, colonies can contain cells in both phenotypes. To determine the distribution of these phenotypes in our different agar conditions, we grew colonies composed of a fluorescent reporter strain. To determine the cellular phenotypic composition of colonies on each agar condition, we performed experiments with a strain containing fluorescent reporters for matrix and motility genes, specifically P*_tapA_*-mCherry for matrix production and P*_hag_*-YFP for motility.^30–33^ We found that the expanding exterior of the 1.5% agar colony was high in the matrix reporter (Fig. 1F), while the exterior of the 0.5% colony is high in the motile reporter (Fig. 1G), consistent with the hypothesis that expansion of biofilms is dominated by matrix-producing cells and expansion of swarms is dominated by motile cells. However, both colonies contained cells in each phenotype. Given that each colony contained cells in both phenotypes, and the two cellular phenotypes are known to drive different physical expansion mechanisms, we wanted to measure motion at the cellular level within both 0.5% and 1.5% agar colonies.

**Figure 1:**
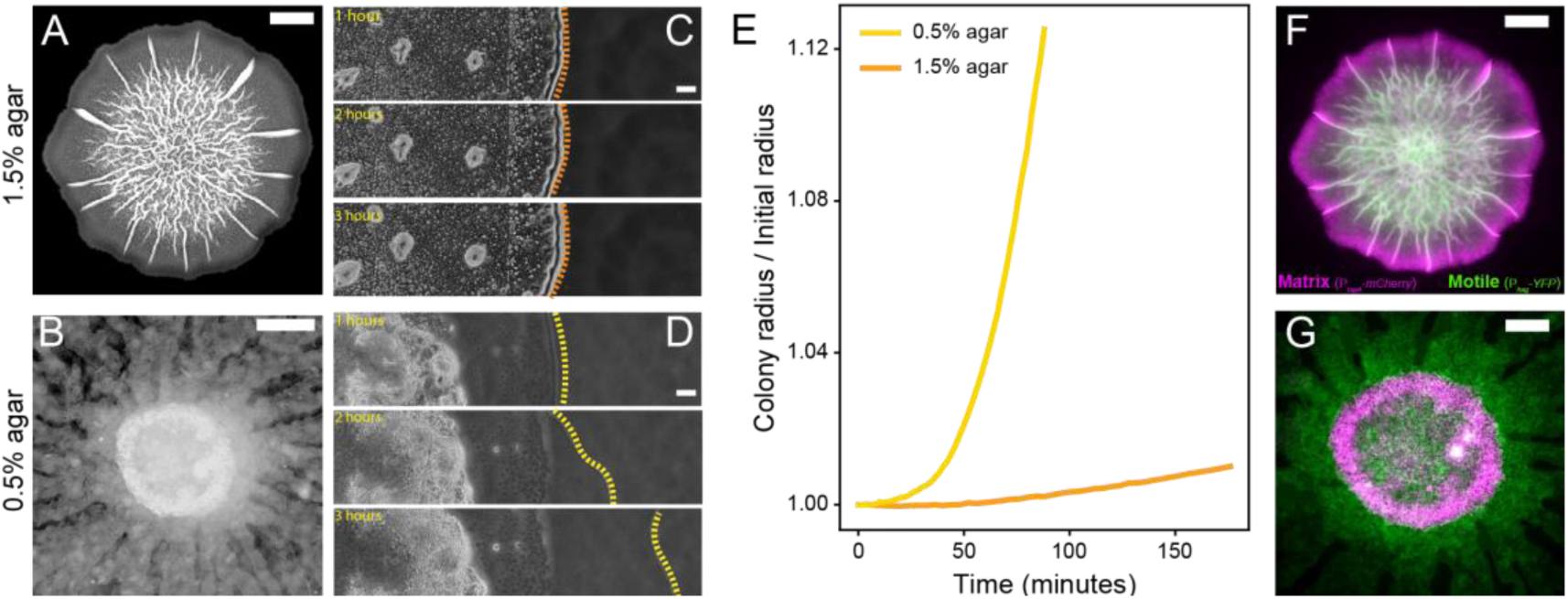
*B. subtilis* displays colony- and cell-level phenotypic variation across environments. **A-B**: Stereoscope images of *B. subtilis* colonies grown on 1.5% and 0.5% agar, respectively. Image A was taken at 24 hours and B at 6 hours. **C-D**: The expanding edge of 1.5% and 0.5% agar colonies, respectively. **E**. Radius over time for 0.5% and 1.5% colonies scaled by the initial inoculum radius. **F,G**: Fluorescence images of colonies from A and B in a reporter strain for matrix production (magenta) and motility (green),

### Bead tracking within colonies reveals cellular level motion

We wanted to track motion at the cellular level within colonies. Tracking individual cells in dense colonies is technically challenging and nearly impossible over large spatial and temporal scales.^34^ To examine cell-level motion during colony growth, we implemented a fluorescent bead tracking assay, similar to others that have been used to probe motion within bacterial colonies.^16,17^ We first deposited a suspension of 1 µm-diameter fluorescent beads (ThermoFisher Scientific; Materials and Methods) onto our agar substrate followed by inoculation of cells in liquid culture (Fig. 2A, Materials and Methods for culture conditions) . This protocol resulted in a dense initial colony of cells with nearly cell-sized beads interspersed (Fig. 2B). Once colonies begin to grow, cells push beads due to growth or motility. By imaging beads at regular time intervals, we can track their trajectories during colony growth. With rapid imaging, we can track bead motion on fast (∼10 s of milliseconds) time scales (Fig. 2C,D,E). Using a particle tracking algorithm, we extract trajectories of many beads within a colony, as illustrated by the isolated bead trajectories in Fig. 2F,G^35^ (Supplementary Video 1). Because we track beads and not cells themselves, our data do not represent the trajectories of cells during colony growth, but rather report passively on motion at the cellular scale within a large colony due to cell activity and growth.

**Figure 2:**
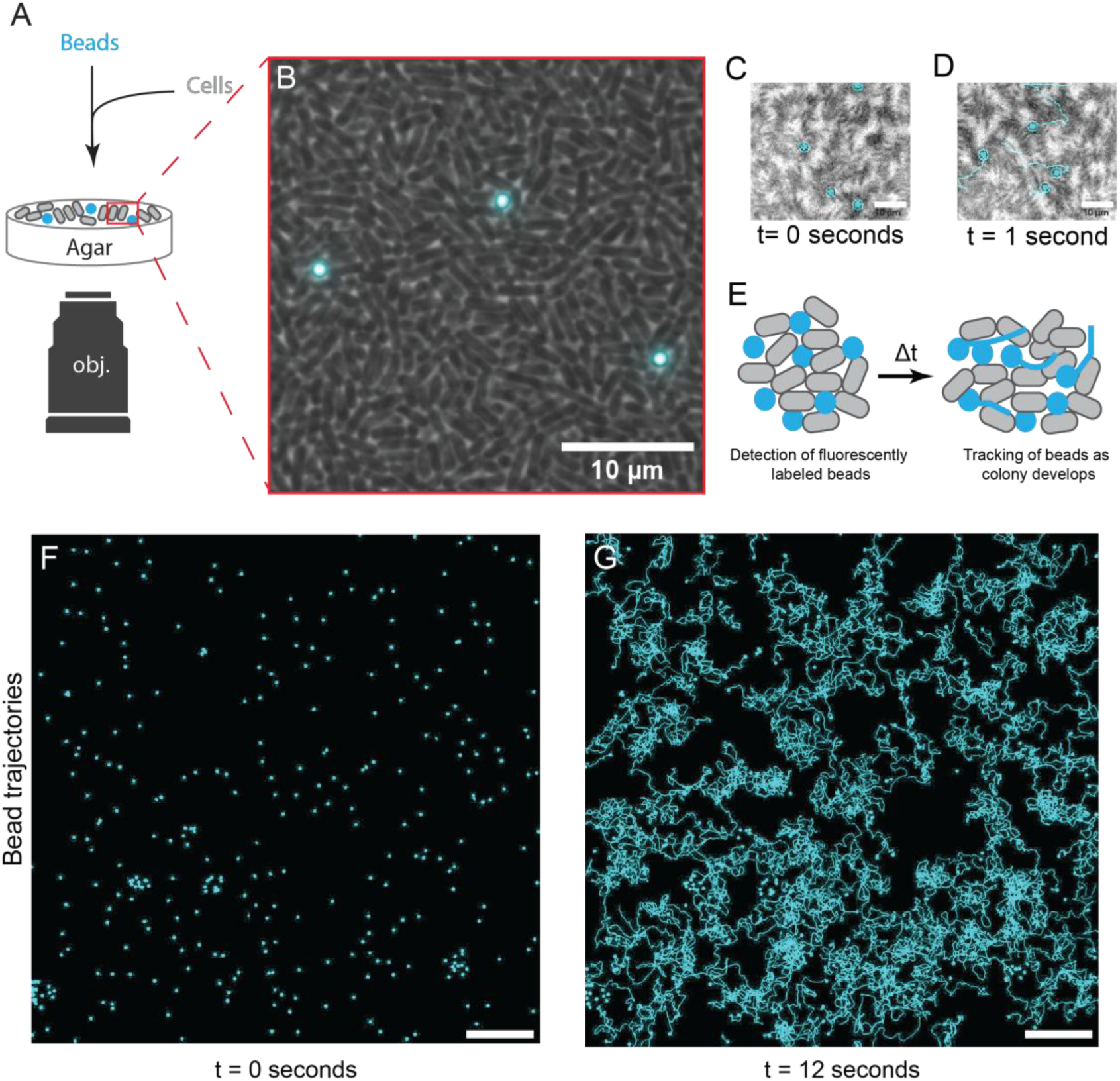
Bead assay reveals cellular level motion in bacterial colonies **A**: Schematic of microscopic setup. **B**: Image of *B. subtilis* cells (gray) and 1 µm fluorescent beads (cyan), scale bar 10 µm. **C-D**: Detection of bead motion between 0 and 1 second. **E**: Schematic of beads with cells for detection and tracking over time. **F-G**: Images of bead fluorescent channel showing detection and motion of beads over 12 seconds of motion (cyan represents bead position and tracked trajectories), scale bare 100 µm.

### Bead-tracking in 0.5% agar colonies reveals bimodal cell-scale motion in swarms

Cells in *B. subtilis* swarms can move rapidly.^36^ For that reason, we tracked beads at high frame rate (∼50 frames per second) to probe motion within these colonies (Fig. 3A). We observed that some beads moved rapidly within swarms, while others moved little or appeared stationary over imaging windows of several seconds (Supplemental Video 2). We computed the average speed over 3-second imaging intervals for these beads and found a bimodal distribution of speeds (Fig. 3B). The fast bead population exhibited speeds on the same order of magnitude as swarm cells or isolated motile cells, (∼10 μm/s).^37^ The slow population moved roughly two orders of magnitude slower (∼0.1 μm/s). Because our high frame-rate imaging necessitated very short imaging intervals, we could not be sure if our observed slow beads were stationary over the longer time-scale of colony growth (Fig. 1E) or were just moving slowly. Our measurements would not be able to precisely determine the character of slow bead trajectories because the slow beads did not exhibit sufficient displacement over the course of the fast-imaging experiments.^38,39^ Moreover, the computed speeds for slow beads were likely an overestimate as small jitters or displacements over such small inter-frame time intervals would make the computed speed artificially high. For that reason, we performed a second set of tracking experiments with much slower frame rate (1 image per minute) to track beads over long periods of time (Fig. 3C). In these slow tracking experiments, we would not be able to follow the fast-moving beads of our high frame-rate experiments (their trajectories would be under-sampled, Fig. S6), but we could follow slow beads for many hours of growth and over the full spatial extent of a colony.

**Figure 3:**
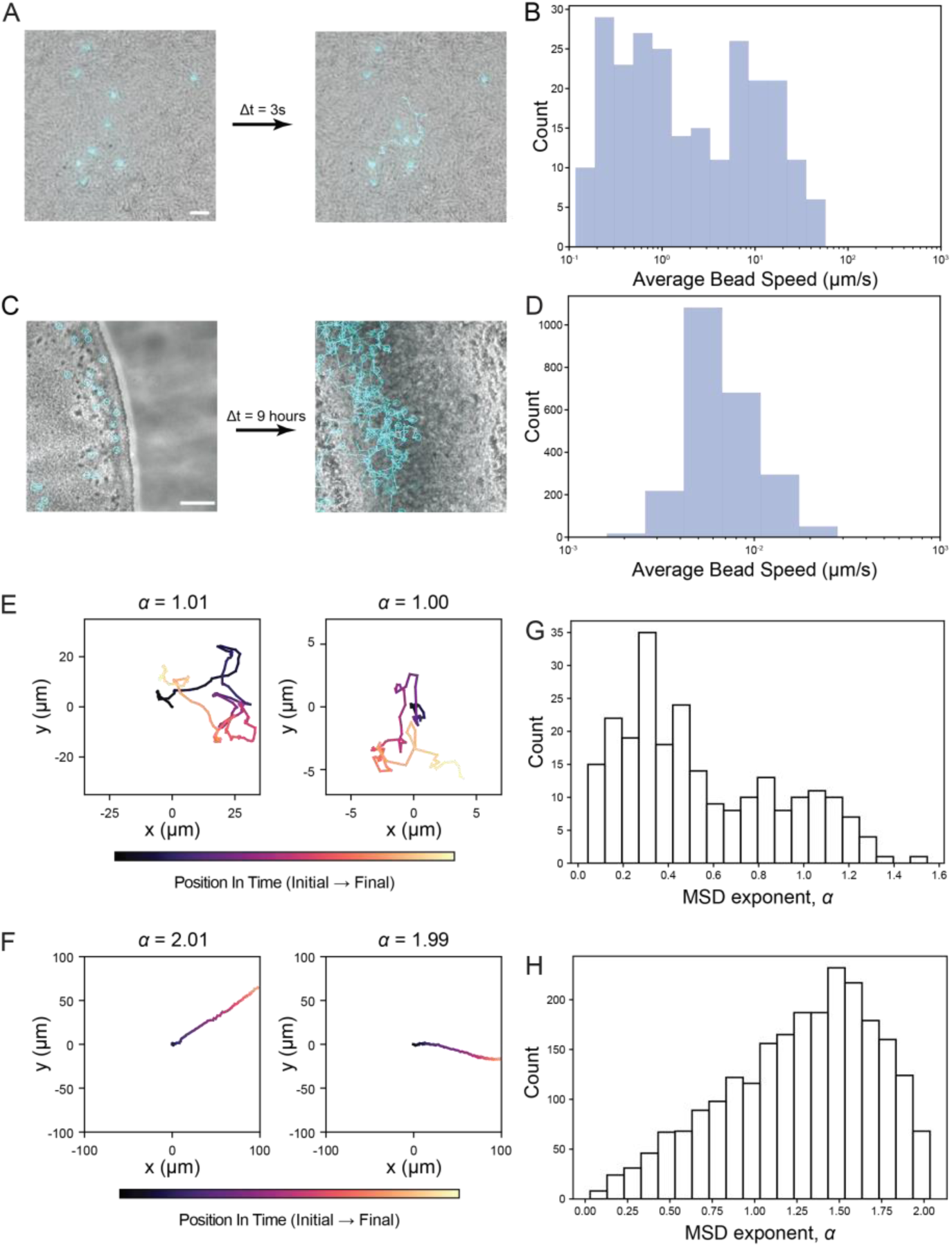
Bead motion is bimodal within swarms. **A**: Fluorescent beads (cyan) embedded in colonies on 0.5% agar after 3 seconds of motion. **B**: Histogram of measured average bead speeds over 3-second intervals. **C**: Fluorescent bead motion over 9 hours and (**D**) average bead velocities. **E**: Sample bead trajectories from fast imaging experiments. **F**: Sample trajectories from slow imaging experiments. **G-H**: Histogram of MSD exponents, α, for fast and slow imaging experiments, respectively.

Extended time lapse measurements revealed that the slow beads were not stationary. They exhibited a monomodal distribution of average velocities centered at 0.005 μm/s (Fig. 3D). Over many hours of colony growth, beads moved substantially, some over 100 µm. The shapes of the trajectories we measured at short (Fig. 3A) and extended (Fig. 3C) imaging time were often qualitatively different. Many fast frame-rate bead trajectories showed many changes in direction (Fig. 3E), while slow frame-rate trajectories were characterized by smooth lines (Fig. 3F). To quantify bead motion in 0.5% agar swarms, we computed the mean squared displacement (MSD) exponent (α) for each bead trajectory. For beads moving ballistically over time, α ≈ 2, i.e. beads tend to move linearly in time and trajectories are directed (see Fig. S4, left for examples). For beads moving with diffusive, Brownian motion, α ≈ 1, and for beads moving sub-diffusively, α ≈ 0 (Supplemental 4). On short time scales (50 fps imaging), we confined our measurement of α to the fast bead population because the population of slow beads did not move enough during our imaging timescale to determine α or the mean speed reliably (Materials and Methods).^38,39^ The fast-moving beads exhibited a bimodal distribution of α values centered at α = 0.31 and α = 1.05, showing a population of sub-diffusive beads (which could be adhered to the substrate) and a population of diffusing beads within the swarm (Fig. 3G). The tracked beads in our 12-hour, low frame-rate time-lapse experiments had a broad distribution of α values peaked at α = 1.48. Our time-lapse experiments have shown that on 0.5% agar substrates, colony-embedded beads move in two different regimes: one population moves rapidly with Brownian motion-like trajectories; the other population moves orders of magnitude more slowly with more ballistic character (Fig. S1).

### Biofilms have heterogenous bead motion during colony growth

Due to the differences in expansion and phenotypic composition of 0.5% and 1.5% MSgg agar colonies (Fig. 1), we hypothesized that bead motion would be qualitatively different between these conditions. We tracked beads in *B. subtilis* biofilms grown on 1.5% agar MSgg (Fig. 4A). As in 0.5% colonies, we first tracked beads at a high frame rate for short time intervals. Unlike in 0.5% swarm conditions, we observed no population of rapidly moving beads. Instead, at high frame rate, we observed a monomodal distribution of bead velocities (Fig. 4B). To track the motion of these slow beads over the course of colony growth, we performed time-lapse imaging experiments of entire 1.5% agar colonies for 12 hours. Over this time scale, we observed a monomodal distribution of average bead velocities centered at v = 0.004 μm/s (Fig. 4C). We computed the MSD exponents for the trajectories of this bead population and found a broad spectrum between 0 and 2 centered at α = 0.820. These results show that over the course of initial colony growth on 1.5% agar, *B. subtilis* biofilms drive strictly slow bead motion, but with highly heterogeneous trajectories (Fig 4D). In addition to trajectories with ballistic and diffusive character, we also resolved beads that exhibited sub-diffusive motion (Fig 4E), characteristic of being trapped within boundaries.^40^ In our swarm and biofilm bead-tracking experiments, we observed slow-moving beads with highly heterogeneous trajectories in both conditions, but only in swarms did we observe a population of fast, diffusing beads. We wanted to know whether this emergent motion in *B. subtilis* colonies arises from the presence of motile or matrix producing cells in *B. subtilis* biofilms and swarms (Fig. 1F,G).

**Figure 4:**
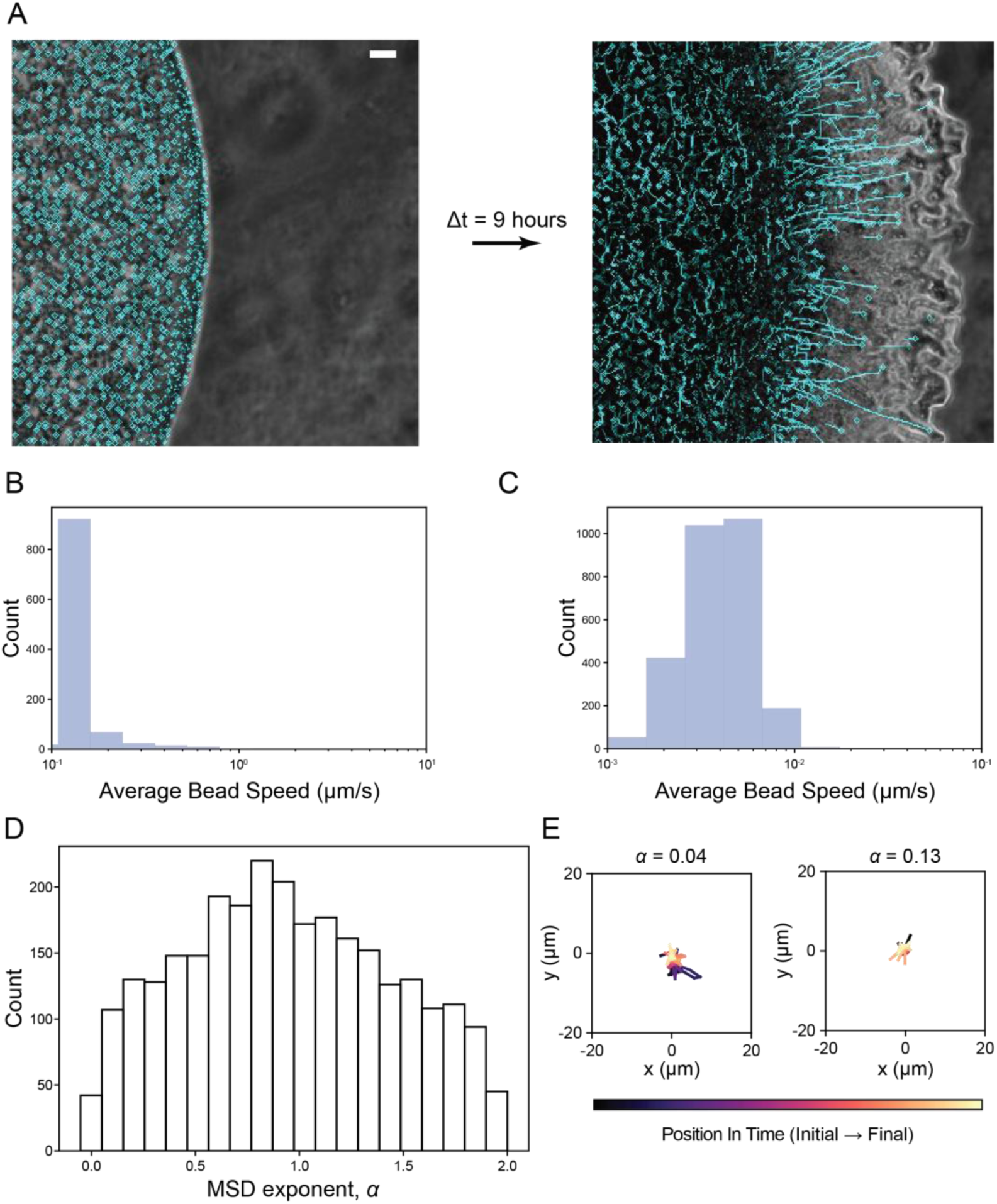
Bead motion is slow and heterogeneous in 1.5% agar colonies. **A**: Fluorescent beads (cyan) embedded in colonies on 1.5% agar after 9 hours of motion, scale bar 100 µm. **B**: Histogram of average bead speeds for fast imaging. **C**: Histogram of bead speeds for slow imaging. **D**: Histogram of MSD exponents for slow imaging. **E**: Sample trajectories of sub-diffusive beads.

### Motile- and *Matrix-only* mutants exhibit distinctive physical properties throughout colony formation

We hypothesized that the different bead trajectories we observe in each agar condition were associated with the distinct cell phenotypes that dominate the colonies, specifically motile cells in swarms and matrix-producers in biofilms. Isolating the effect of each is difficult because swarms and biofilms each contain cells in both phenotypic states (Fig. 1F,G). To test this hypothesis, we performed bead-tracking experiments with regulatory mutants of *B. subtilis* that are locked into motile and matrix phenotypes respectively. A feedback loop involving the transcriptional repressor *sinR* and its allosteric regulator *sinI* regulates motility and matrix production in *B. subtilis*: *sinR* represses matrix production genes and the SinI protein binds to SinR, inhibiting its repressive activity.^31,32,41,42^ Consequently, in a *sinI* knockout strain (Δ*sinI*, Materials and Methods), matrix production is repressed and all cells are motile (Fig. 5A); in a *sinR* knockout strain (Δ*sinR*, Materials and Methods), all cells produce matrix (Fig. 5C). Δ*sinI* colonies form rapidly expanding, smooth swarms on 0.5% agar (Fig. 5B), while Δ*sinR* colonies form highly robust biofilms on 1.5% agar. We tracked bead motion in colonies composed of these two mutant strains on both 0.5% and 1.5% agar MSgg to isolate the effect of each cellular phenotype in each condition (Fig 5E-L). On 0.5% agar substrates, we observed a monomodal distribution of bead velocities in fast frame-rate movies of Δ*sinI* that was centered at v = 95.4 μm/s (Fig. 5E, see Supplemental Video 3). This contrasts WT, which under the same condition had a population of slower-moving beads, which moved ballistically over the course of colony growth (Fig. 3). We take this as evidence that motile cells alone cannot account for the slow, directed bead motion that arose in WT swarms. Clusters of matrix-producing cells may drive the slow bead motion we observed in 0.5% agar WT swarms. The rapidly moving beads in 0.5% Δ*sinI* had a monomodal α distribution centered at a super-diffusive value of α = 1.49 (Fig. 5F). These α values are higher than those of the diffusive, fast-moving beads in WT (Fig. 3C, Supplemental 5). The trajectories of beads in 0.5% Δ*sinI* colonies had diffusive character punctuated by periods of more directed motion (Supplemental Movie 1). On 1.5% agar substrates, Δ*sinI* colonies showed similar velocity and α distributions to WT (Fig. 5G,H). Beads moved slowly over the course of 12 hours of colony growth, with a broad α distribution centered near α = 1.

**Figure 5:**
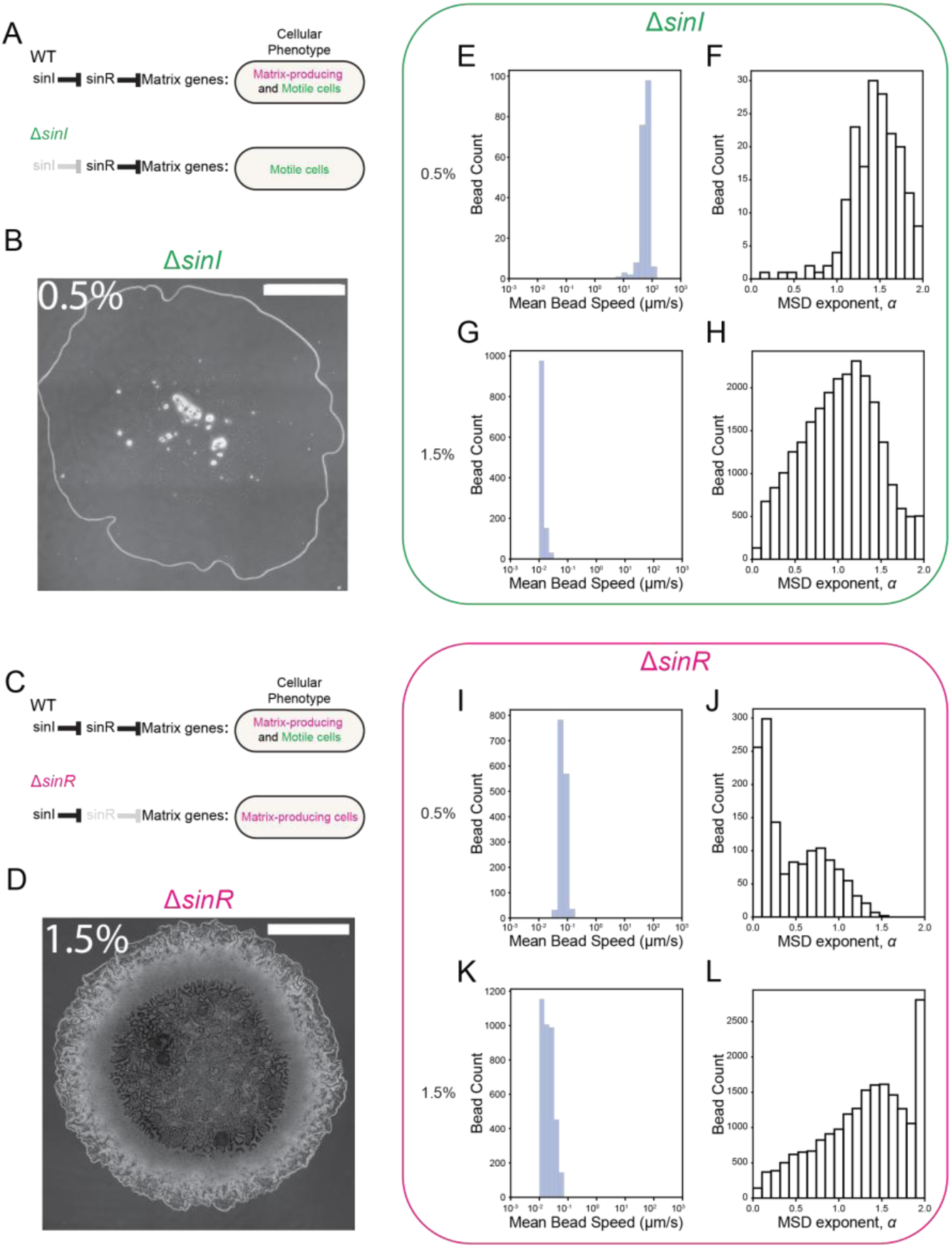
Specific cellular phenotypes are associated with different regimes of motion. **A**: Simplified schematic of gene regulatory network illustrating Δ*sinI*. **B**: Microscope image of Δ*sinI* strain grown on 0.5% agar, scale bar 1000 µm. **C**: Simplified schematic of gene regulatory network illustrating Δ*sinR*. **D**: Microscope image of Δ*sinR* strain grown on 1.5% agar, scale bar 1000 µm. **E**: Mean bead speed and (**F**) MSD exponents for Δ*sinI* on 0.5% agar. **G**: Mean bead speed and (**H**) MSD exponents for Δ*sinI* on 1.5% agar. **I**: Mean bead speed and (**J**) MSD exponents for Δ*sinR* on 0.5% agar. **K**: Mean bead speed and (**L**) MSD exponents for Δ*sinR* on 1.5% agar.

Δ*sinR* colonies exhibited little growth on 0.5% substrates, causing little bead motion. The motion we did observe was low in average velocity and sub-diffusive (Fig 5I,J). This may be due to the inability of Δ*sinR* cells to exhibit flagellar motility, which may be necessary to expand over a soft surface like 0.5% agar. On 1.5% agar substrates, however, Δ*sinR* colonies exhibited a similar distribution of average bead velocities to WT (Fig. 5K). Δ*sinR* also showed a broad distribution of MSD exponents, but with some differences from WT under the same condition: its peak value was slightly higher and it had a prominent second peak at α = 2, showing strong ballistic motion driven by expanding matrix-producers (Fig. 5L). Our results from Δ*sinI* suggest that, within swarms, motile cells drive diffusive bead motion and that slow bead motion in WT swarms is likely due to the presence of matrix-producing clusters that emerge during colony growth. The latter conclusion is supported by the lack of a slow, ballistic bead population in 0.5% Δ*sinI* experiments. In Δ*sinR* experiments, the presence of a prominent α = 2 peak supports the conclusion that matrix-producing cells drive more ballistic motion in biofilms, likely due to strongly radially directed colony expansion.

## Discussion

Our findings reveal distinct patterns of cell-scale motion in *B. subtilis* swarms and biofilms. Specifically, motile cells in swarms exhibit diffusive motion, characterized by non-directional, random trajectories, whereas the presence of matrix-producing cells favors more ballistic motion in both biofilms and swarms. This distinction underscores the fundamentally different physical and regulatory states associated with these cell types, and the results show that heterogeneous cell phenotypes contribute to qualitatively different physics within the same colony. Understanding the growth and phenotypic composition of these colonies will require models that consider the properties of both cell types.

The diffusive motion observed in motile cells is consistent with previously established behaviors of flagellated cells navigating their environment through ‘run-and-tumble’ dynamics. Their stochastic trajectories may allow cells to sample a variety of environmental niches before committing to sessility or differentiation. This is in line with previous models suggesting that motile cells maintain a high degree of phenotypic plasticity and environmental responsiveness.^43^ Conversely, the ballistic motion of matrix-producing cells may reflect directed expansion as matrix cells spread outward due to growth and mechanical pushing.

These observations also invite further mechanistic questions into how regulatory networks involving *SinR* and *SinI* contribute to the emergence of these distinct motion profiles, for example whether cells switch phenotypes and establish different regimes of motion within colonies. Furthermore, we do not know how the physical properties of the extracellular matrix facilitate or constrain cell motion. Future work utilizing spatial analysis of mutant strains could illuminate the correlative relationships between gene expression programs, cell fate decisions, and movement dynamics.

## Materials and Methods

### Bacillus subtilis

We use strain NCIB3610 of the bacterial species *Bacillus subtilis*. All mutants were transformed using NCIB3610 as parent strain and verified via inserting an antibiotic resistance into the genome and selecting colonies that were positively resistant, as well as sequencing. (Refer to Table 1.1 for strain descriptions.)

**Table 1.1.** Strains of *Bacillus subtilis*.

| **Strain** | **Genotype** | **Source** |
| --- | --- | --- |
| Wild type | NCIB3610 pBS32 (comI<br>Q12L) plasmid point<br>mutation | BGSC (Irnov & Winkler,<br>2010)<sup>44</sup> |
| *ΔsinI* | NCIB3610 pBS32(comI<br>Q12L) ΔsinI::mls | This study |
| *ΔsinR* | NCIB3610 pBS32(comI<br>Q12L) ΔsinR::mls | This study |

### Fluorescent beads

Beads for motion tracking were 1.0µm diameter FluoSpheres™ Carboxylate-Modified Microspheres (ThermoFisher Scientific).

### Colony growth on MSgg agar plates

All *B. subtilis* strains were grown on Luria Broth (LB) plates overnight and cultured in liquid LB on the day of imaging. Cells were grown in liquid LB medium at 30°C for approximately 2.5 – 3 hours, until reaching late exponential phase, measured by an OD of 0.70 to 0.85. The cultures were then diluted to a ratio of 1:20 and subsequently resuspended in liquid MSgg medium. The cultures were grown for 3 hours reaching an OD of 0.80 to 1.00, indicating that the cells are in their logarithmic phase of growth, and were ready for plating. The MSgg Agar plates were then inoculated with 1µL of fluorescent beads followed by 1µL of matured cultures. Plates were then covered and Parafilm was applied to minimize the loss of moisture to the air.

The MSgg media was composed of 5 mM potassium phosphate buffer (pH 7.0), 100 mM MOPS buffer (pH 7.0, adjusted using NaOH), 2 mM MgCl2, 700 µM CaCl2, 50 µM MnCl2, 100 µM FeCl3, 1 µM ZnCl2, 2 µM thiamine HCl, 0.5% (v/v) glycerol and 0.5% (w/v) monosodium glutamate. The MSgg agar plates include a 0.5% or 1.5 concentration of agar, measured in weight per volume, in the MSgg media.

### Microscopy and image analysis

Time lapse images were taken using a 10X objective 0.3 NA objective on an Olympus IX83 epifluorescence microscope. For time lapse imaging of whole colony imaging, images were taken every 1 minute for up to12 hours. Images of strains including fluorescent reporters were taken every 3 minutes in phase contrast and fluorescence channels, mCherry, and YFP channels for up to 12 hours. Fluorescent beads were imaged in mCherry or DAPI fluorescent channels. Mutant strain images, for *ΔsinI* and *ΔsinR* colonies, were taken in phase contrast. Fast imaging occurred for intervals of 3-10 seconds and collected images every 0.02 seconds. This is equivalent to imaging at 50 frames per second. Still images of beads localized with cells were taken using a 20x objective 0.3 NA on an Olympus IX83 epifluorescence microscope. Still images of full colonies were taken using an Olympus MVX10 Macro Zoom Scope at 0.63x – 1.0x magnification range. All images were collected in Olympus CellSens software, then exported and processed using ImageJ. Post processing and analysis occurred using Mathematica and Python scripts.

### Tracking of fluorescent beads

The ImageJ plug-in MOSAICs particle-tracker was used in tracking the trajectories of fluorescent particles.^35^ Diffusion coefficients and linear fits of each particles trajectories were computed by MOSAIC as well and used directly in plotting the results. The single particle tracker outputs trajectory data over the course of the imaging period, from which it generates a linear fit of the mean squared displacement evolution of each particle. The linear fit was then used to derive the MSD exponent [<r^2^(t)> = k t^α^] used in generating histograms. The average speed of each particle was also provided by MOSAIC, in units of [m/frame] which was then converted to [m/s] and the histogram was plotted. Note: this method is not tracking cells, but rather the passive cellular scale motion arising in the colony.

---

**Fig. S1:**
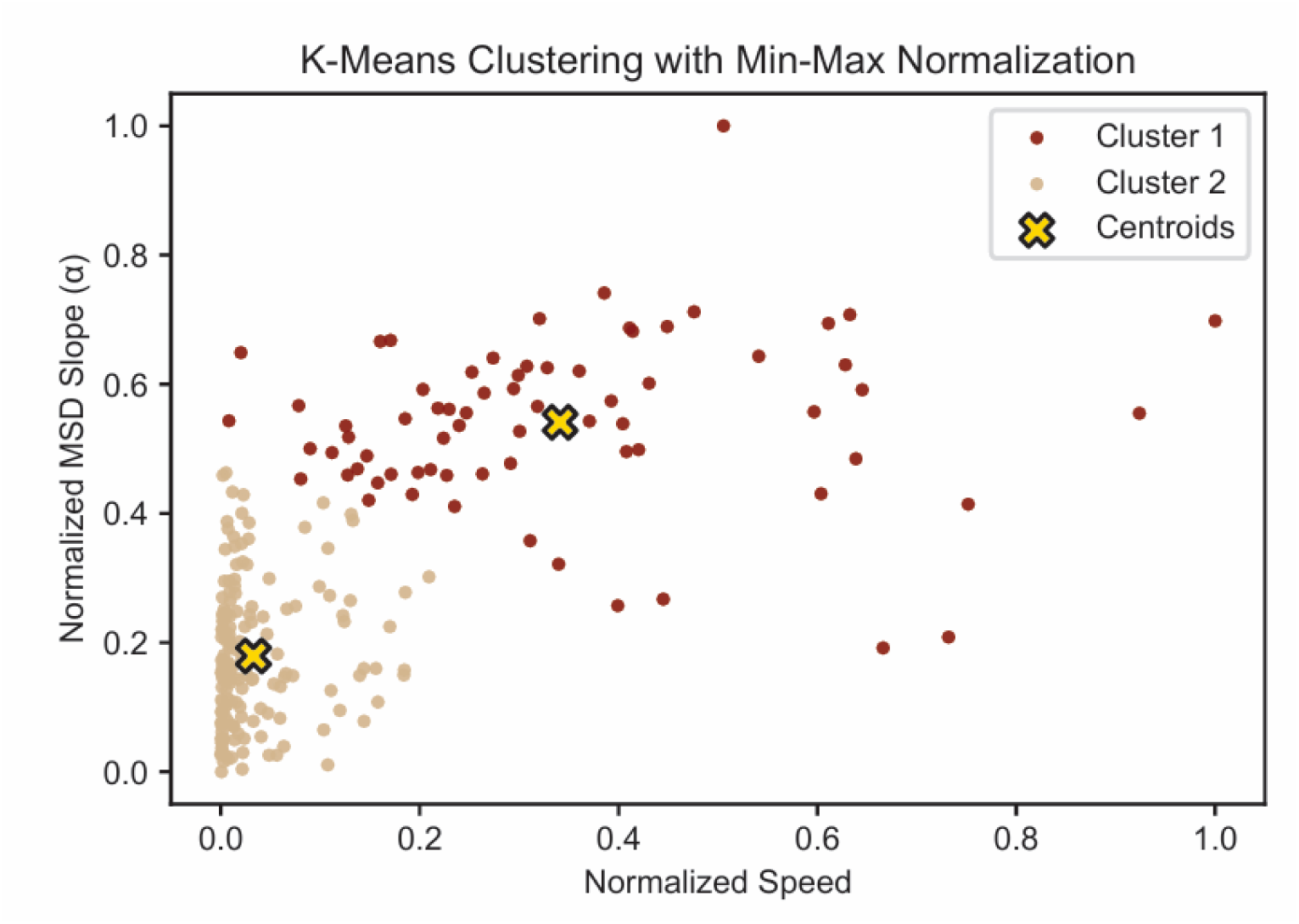
K-Means clustering identifies distinct populations of bead motion in colonies grown on 0.5% agar during fast imaging in mean-normalized MSD exponent and speed values of Wild Type.

**Fig S2:**
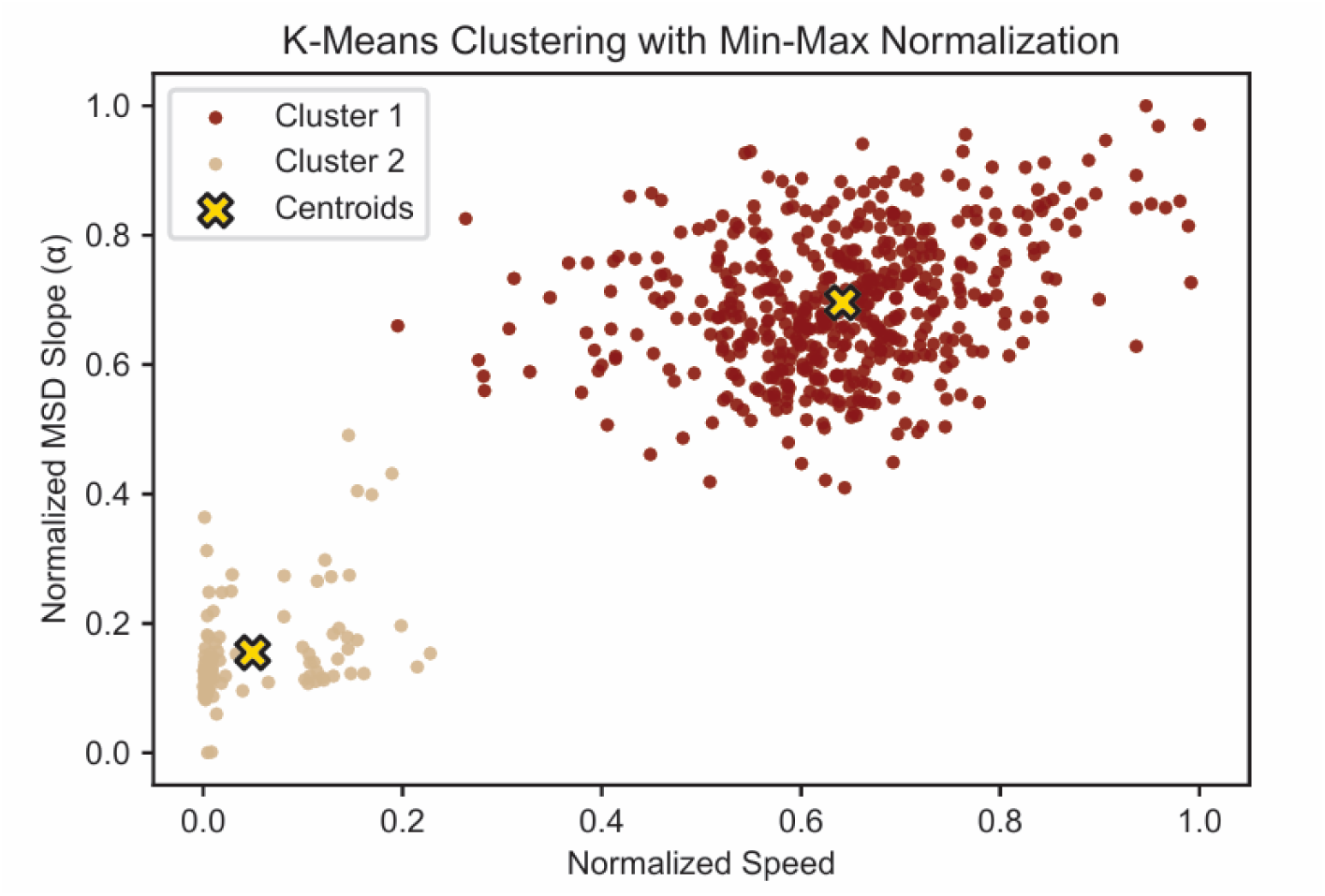
K-Means clustering identifies distinct populations of bead motion in colonies grown on 0.5% agar during fast imaging in mean-normalized MSD exponent and speed values for Δ*sinI*.

**Fig S3:**
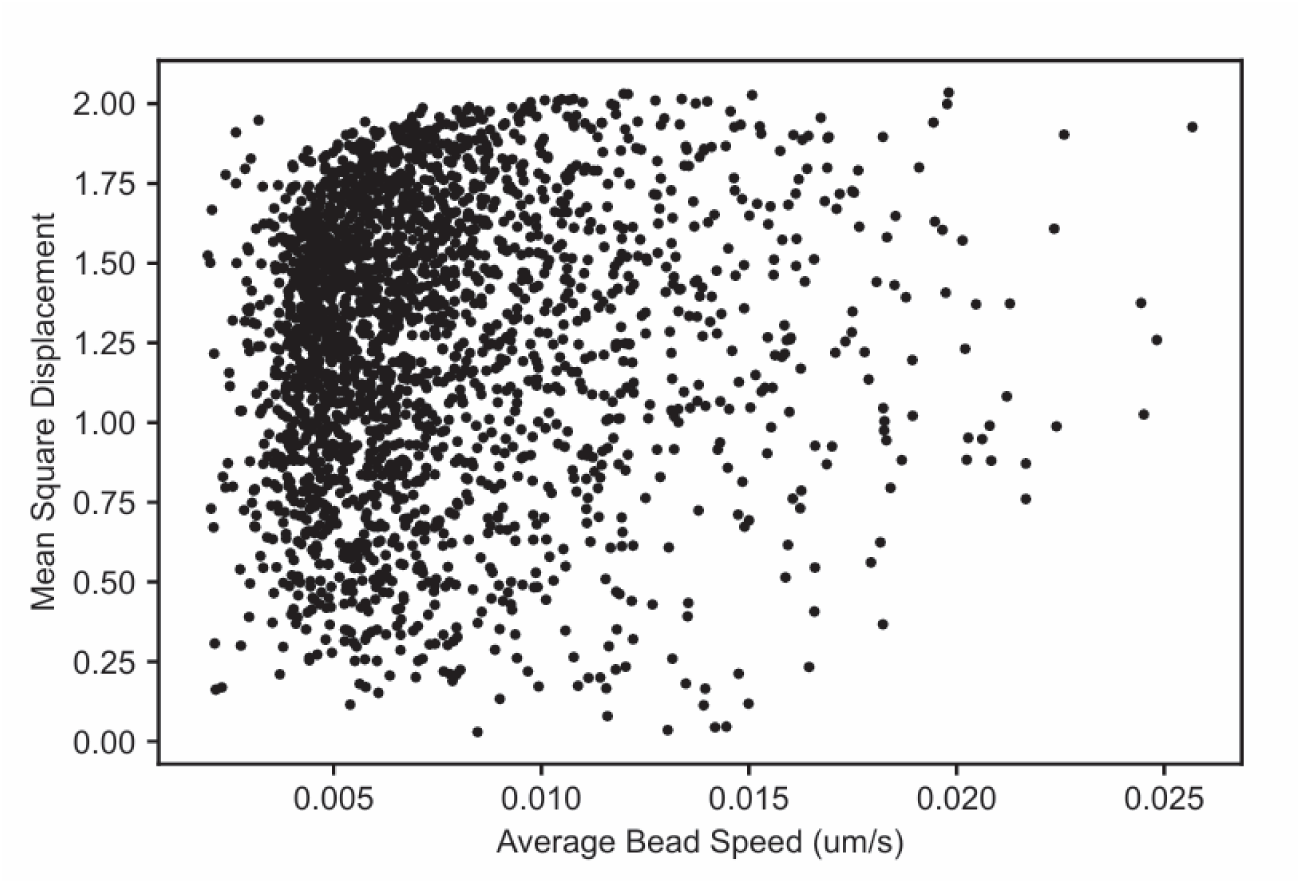
Scatterplot of average bead speed and MSD exponent values for Wild Type in whole colonies grown on 0.5% agar during slow imaging.

**Fig. S4:**
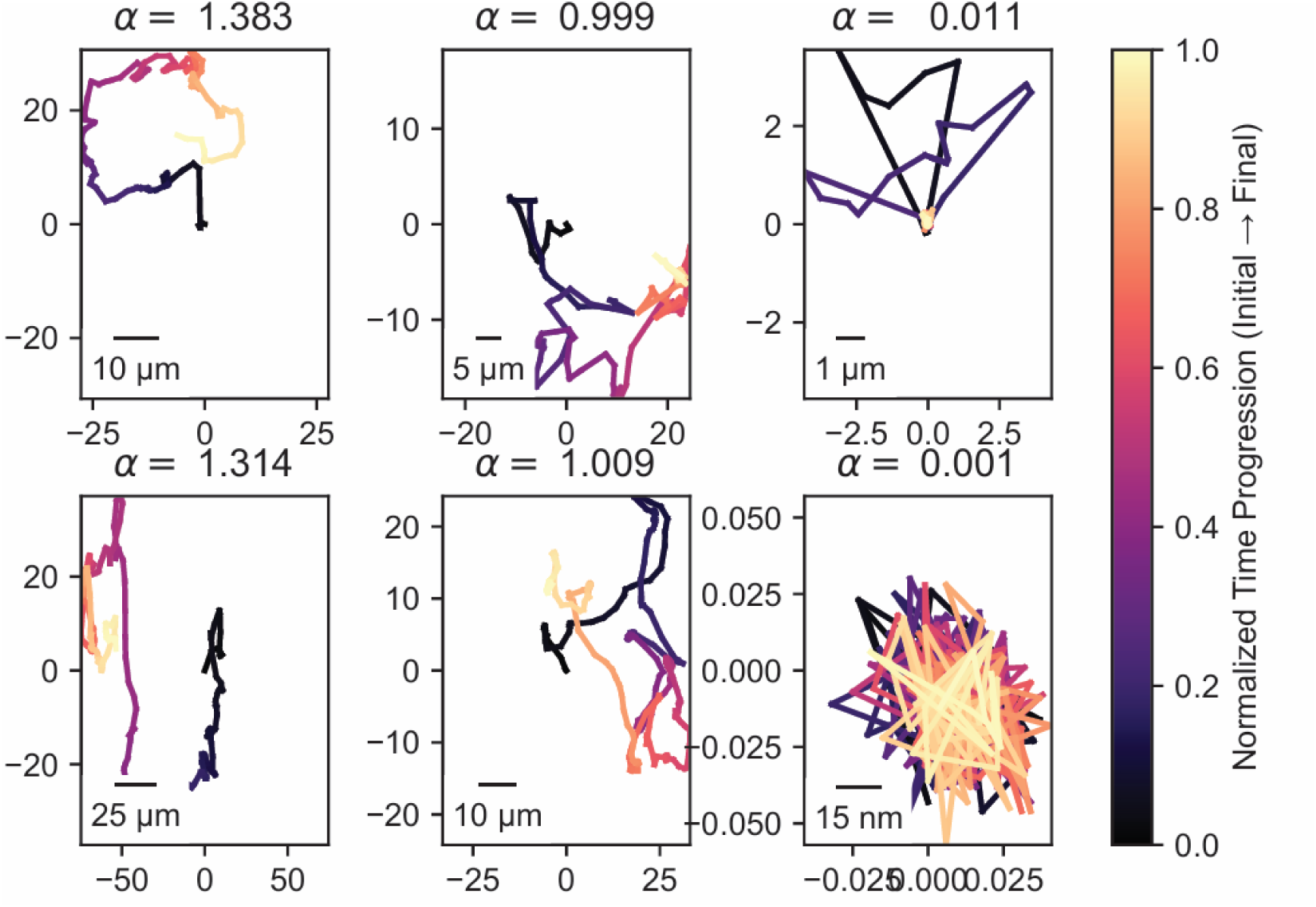
Full examples of MSD trajectories across quantities of alpha. Showcases trajectories 1 < *α* < 2 (left column) as well as trajectories at higher spatial resolutions.

**Fig. S5:**
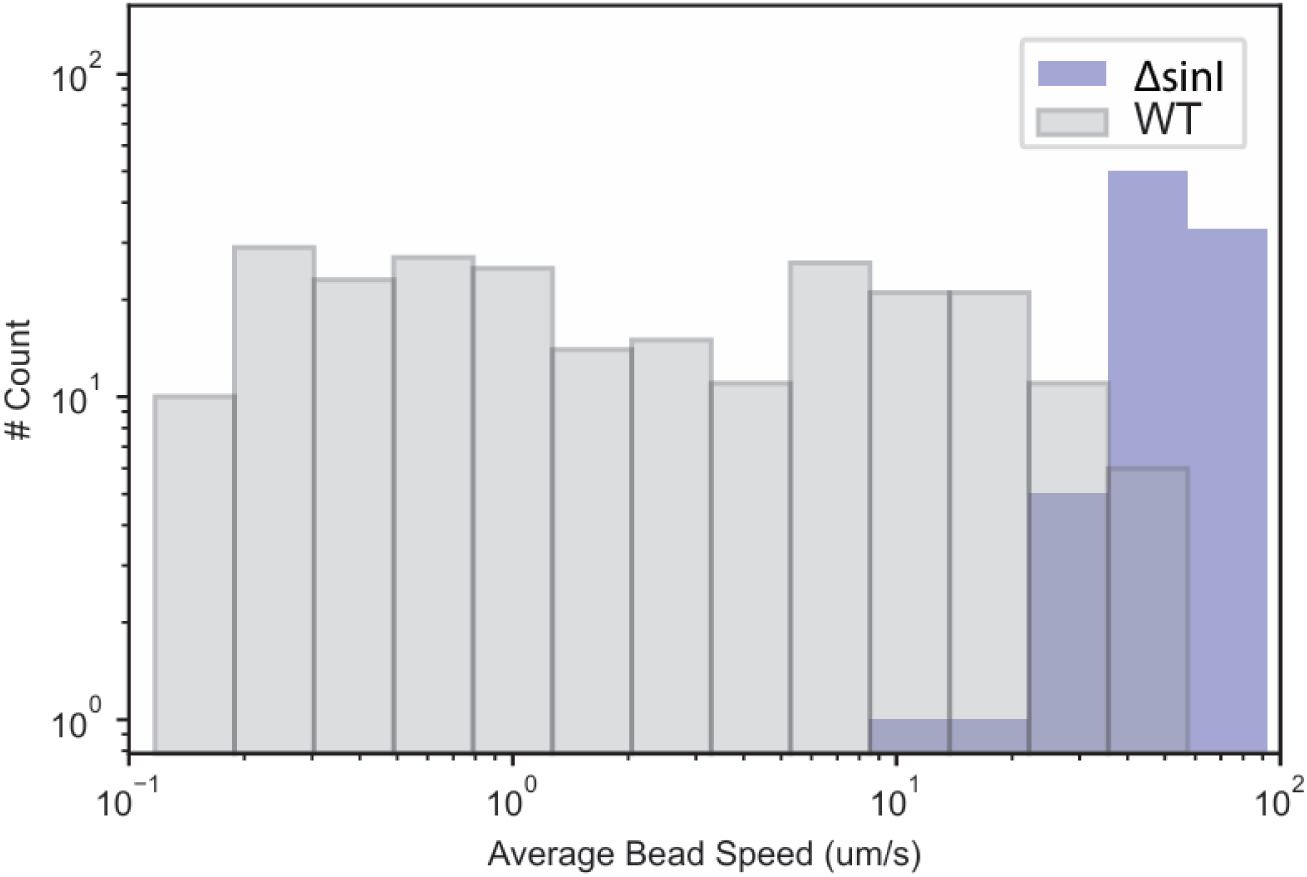
Speed histogram comparing colonies grown on 0.5% agar: Δ*sinI* (blue) and WT (grey) during fast imaging modality.

**Fig. S6:**
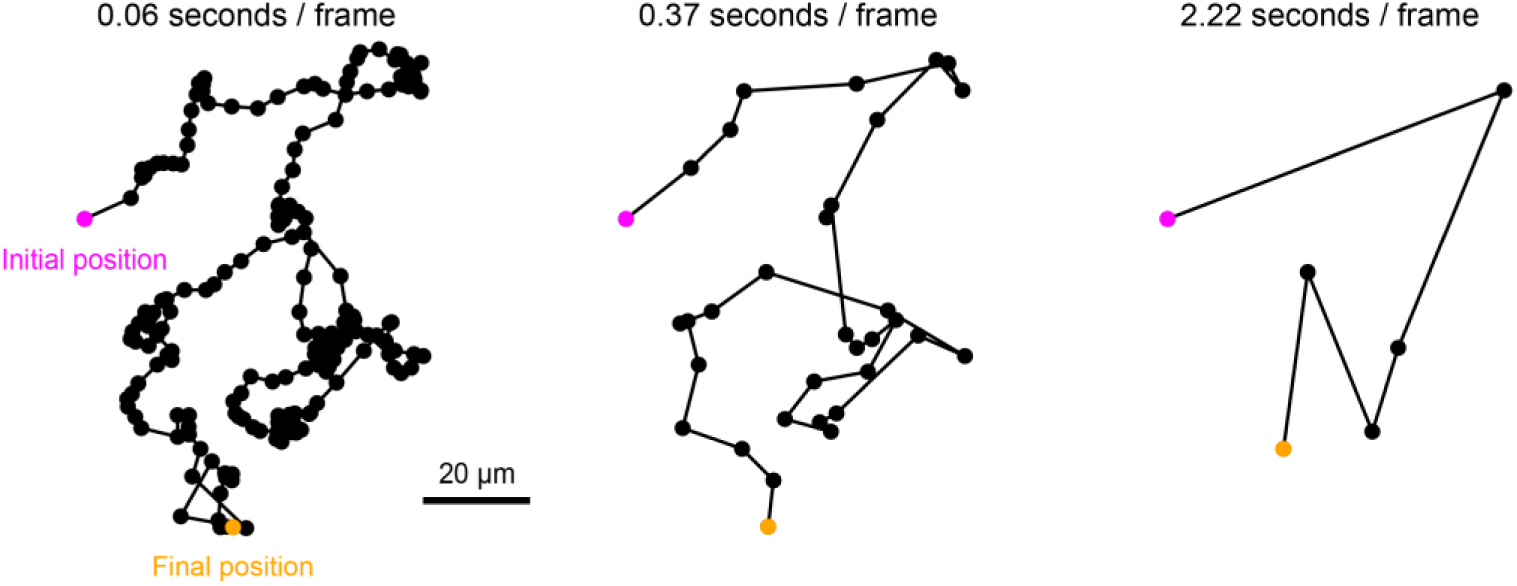
A bead trajectory with increasingly slower sampling. Exemplifying the characteristic difference between trajectories obtained from slower imaging modalities.

## Videos

**Supplemental Video 1.** Example of a tracked bead fluorescence movie (Δ*sinI* on 0.5% agar). Beads are white. Tracked trajectories are colored.

**Supplemental Video 2.** Phase contrast movie of 1 µm beads (dark) being pushed by a WT *B. subtilis* swarm on 0.5% agar MSgg.

**Supplemental Video 3.** Phase contrast movie of 1 µm beads (dark) being pushed by a Δ*sinI B. subtilis* swarm on 0.5% agar MSgg.

## References

1. Köhler, T., Curty, L. K., Barja, F., Van Delden, C. & Pechère, J.-C. Swarming of *Pseudomonas aeruginosa* Is Dependent on Cell-to-Cell Signaling and Requires Flagella and Pili. J. Bacteriol. 182, 5990–5996 (2000).

2. Kearns, D. B. A field guide to bacterial swarming motility. Nat. Rev. Microbiol. 8, 634–644 (2010).

3. Vlamakis, H., Aguilar, C., Losick, R. & Kolter, R. Control of cell fate by the formation of an architecturally complex bacterial community. Genes Dev. 22, 945–953 (2008).

4. Shemesh, M. & Chai, Y. A Combination of Glycerol and Manganese Promotes Biofilm Formation in Bacillus subtilis via Histidine Kinase KinD Signaling. J. Bacteriol. 195, 2747–2754 (2013).

5. Sauer, K., Camper, A. K., Ehrlich, G. D., Costerton, J. W. & Davies, D. G. *Pseudomonas aeruginosa* Displays Multiple Phenotypes during Development as a Biofilm. J. Bacteriol. 184, 1140–1154 (2002).

6. Branda, S. S., González-Pastor, J. E., Ben-Yehuda, S., Losick, R. & Kolter, R. Fruiting body formation by *Bacillus subtilis*. Proc. Natl. Acad. Sci. 98, 11621–11626 (2001).

7. Kearns, D. B. & Losick, R. Swarming motility in undomesticated *Bacillus subtilis*. Mol. Microbiol. 49, 581–590 (2003).

8. Matsushita, M. et al. Colony formation in bacteria: experiments and modeling. Biofilms 1, 305–317 (2004).

9. Du, H., Xu, Z., Shrout, J. D. & Alber, M. MULTISCALE MODELING OF *PSEUDOMONAS AERUGINOSA* SWARMING. Math. Models Methods Appl. Sci. 21, 939–954 (2011).

10. Trinschek, S., John, K. & Thiele, U. Modelling of surfactant-driven front instabilities in spreading bacterial colonies. Soft Matter 14, 4464–4476 (2018).

11. Jeckel, H. et al. Learning the space-time phase diagram of bacterial swarm expansion. Proc. Natl. Acad. Sci. 116, 1489–1494 (2019).

12. Srinivasan, S., Kaplan, C. N. & Mahadevan, L. A multiphase theory for spreading microbial swarms and films. eLife 8, e42697 (2019).

13. Yan, J. et al. Mechanical instability and interfacial energy drive biofilm morphogenesis. eLife 8, e43920 (2019).

14. Fei, C. et al. Nonuniform growth and surface friction determine bacterial biofilm morphology on soft substrates. Proc. Natl. Acad. Sci. 117, 7622–7632 (2020).

15. Cont, A., Rossy, T., Al-Mayyah, Z. & Persat, A. Biofilms deform soft surfaces and disrupt epithelia. eLife 9, e56533 (2020).

16. Charlton, S. G. V., Kurz, D. L., Geisel, S., Jimenez-Martinez, J. & Secchi, E. The role of biofilm matrix composition in controlling colony expansion and morphology. Interface Focus 12, 20220035 (2022).

17. Morris, R. J., Stevenson, D., Sukhodub, T., Stanley-Wall, N. R. & MacPhee, C. E. Density and temperature controlled fluid extraction in a bacterial biofilm is determined by poly-γ-glutamic acid production. Npj Biofilms Microbiomes 8, 98 (2022).

18. Guttenplan, S. B. & Kearns, D. B. Regulation of flagellar motility during biofilm formation. FEMS Microbiol. Rev. 37, 849–871 (2013).

19. Worlitzer, V. M. et al. Biophysical aspects underlying the swarm to biofilm transition. Sci. Adv. 8, eabn8152 (2022).

20. Kinsinger, R. F., Shirk, M. C. & Fall, R. Rapid Surface Motility in *Bacillus subtilis* Is Dependent on Extracellular Surfactin and Potassium Ion. J. Bacteriol. 185, 5627–5631 (2003).

21. Kearns, D. B. & Losick, R. Cell population heterogeneity during growth of *Bacillus subtilis*. Genes Dev. 19, 3083–3094 (2005).

22. Norman, T. M., Lord, N. D., Paulsson, J. & Losick, R. Memory and modularity in cell-fate decision making. Nature 503, 481–486 (2013).

23. Srinivasan, S. et al. Matrix Production and Sporulation in Bacillus subtilis Biofilms Localize to Propagating Wave Fronts. Biophys. J. 114, 1490–1498 (2018).

24. Jeckel, H. et al. Simultaneous spatiotemporal transcriptomics and microscopy of Bacillus subtilis swarm development reveal cooperation across generations. Nat. Microbiol. 8, 2378–2391 (2023).

25. Wang, X., Kong, Y., Zhao, H. & Yan, X. Dependence of the *Bacillus subtilis* biofilm expansion rate on phenotypes and the morphology under different growing conditions. Dev. Growth Differ. 61, 431–443 (2019).

26. Mattingly, A. E., Weaver, A. A., Dimkovikj, A. & Shrout, J. D. Assessing Travel Conditions: Environmental and Host Influences on Bacterial Surface Motility. J. Bacteriol. 200, (2018).

27. Worlitzer, V. M. et al. Biophysical aspects underlying the swarm to biofilm transition. Sci. Adv. 8, eabn8152 (2022).

28. Errington, J. Determination of cell fate in Bacillus subtilis. Trends Genet. 12, 31–34 (1996).

29. Vlamakis, H., Chai, Y., Beauregard, P., Losick, R. & Kolter, R. Sticking together: building a biofilm the Bacillus subtilis way. Nat. Rev. Microbiol. 11, 157–168 (2013).

30. Kearns, D. B., Chu, F., Branda, S. S., Kolter, R. & Losick, R. A master regulator for biofilm formation by *Bacillus subtilis*. Mol. Microbiol. 55, 739–749 (2005).

31. Colledge, V. L. et al. Structure and Organisation of SinR, the Master Regulator of Biofilm Formation in Bacillus subtilis. J. Mol. Biol. 411, 597–613 (2011).

32. Kampf, J. et al. Selective Pressure for Biofilm Formation in Bacillus subtilis: Differential Effect of Mutations in the Master Regulator SinR on Bistability. mBio 9, e01464–18 (2018).

33. Niklas, K. J. & Newman, S. A. The many roads to and from multicellularity. J. Exp. Bot. 71, 3247–3253 (2020).

34. Kodakkat, S. et al. Biofilm characterization: Imaging, analysis and considerations. In Methods in Microbiology vol. 54 39–79 (Elsevier, 2024).

35. Bernhardt, N. & Faraldo-Gómez, J. D. MOSAICS: A software suite for analysis of membrane structure and dynamics in simulated trajectories. Biophys. J. 122, 2023–2040 (2023).

36. Jeckel, H. et al. Learning the space-time phase diagram of bacterial swarm expansion. Proc. Natl. Acad. Sci. 116, 1489–1494 (2019).

37. Mendelson, N. H., Bourque, A., Wilkening, K., Anderson, K. R. & Watkins, J. C. Organized Cell Swimming Motions in *Bacillus subtilis* Colonies: Patterns of Short-Lived Whirls and Jets. J. Bacteriol. 181, 600–609 (1999).

38. Kepten, E., Weron, A., Sikora, G., Burnecki, K. & Garini, Y. Guidelines for the Fitting of Anomalous Diffusion Mean Square Displacement Graphs from Single Particle Tracking Experiments. PLOS ONE 10, e0117722 (2015).

39. Kepten, E., Bronshtein, I. & Garini, Y. Improved estimation of anomalous diffusion exponents in single-particle tracking experiments. *Phys*. Rev. E 87, 052713 (2013).

40. Muñoz-Gil, G. et al. Objective comparison of methods to decode anomalous diffusion. Nat. Commun. 12, 6253 (2021).

41. Milton, M. E. et al. The Solution Structures and Interaction of SinR and SinI: Elucidating the Mechanism of Action of the Master Regulator Switch for Biofilm Formation in Bacillus subtilis. J. Mol. Biol. 432, 343–357 (2020).

42. Dannenberg, S., Penning, J., Simm, A. & Klumpp, S. The motility-matrix production switch in *Bacillus subtilis* —a modeling perspective. J. Bacteriol. 206, e00047–23 (2024).

43. Colin, R., Ni, B., Laganenka, L. & Sourjik, V. Multiple functions of flagellar motility and chemotaxis in bacterial physiology. FEMS Microbiol. Rev. 45, fuab038 (2021).

44. Irnov, I. & Winkler, W. C. A regulatory RNA required for antitermination of biofilm and capsular polysaccharide operons in Bacillales: Antitermination of exopolysaccharide genes. Mol. Microbiol. 76, 559–575 (2010).

